# Implicit reward associations impact face processing: Time-resolved evidence from event-related brain potentials and pupil dilations

**DOI:** 10.1101/232538

**Authors:** Wiebke Hammerschmidt, Igor Kagan, Louisa Kulke, Annekathrin Schacht

## Abstract

The present study aimed at investigating whether associated motivational salience causes preferential processing of inherently neutral faces similar to emotional expressions by means of event-related brain potentials (ERPs) and changes of the pupil size. To this aim, neutral faces were implicitly associated with monetary outcome, while participants (N = 44) performed a subliminal face-matching task that ensured performance around chance level and thus an equal proportion of gain, loss, and zero outcomes. Motivational context strongly impacted processing of all – even task-irrelevant – stimuli prior to the target face, indicated by enhanced amplitudes of subsequent ERP components and increased pupil size. In a separate test session, previously associated faces as well as novel faces with emotional expressions were presented within the same task but without motivational context and performance feedback. Most importantly, previously gain-associated faces amplified the LPC, although the individually contingent face-outcome assignments were not made explicit during the learning session. Emotional expressions impacted the N170 and EPN components. Modulations of the pupil size were absent in both motivationally-associated and emotional conditions. Our findings demonstrate that neural representations of neutral stimuli can acquire increased salience via implicit learning, with an advantage for gain over loss associations.

## Introduction

To support adaptive behavior in complex environments, the human brain developed efficient selection mechanisms that bias perception in favor of salient information. In order to address the variety of different sources of salience, conventional attention theories focusing on goal- and salience-driven attention mechanisms (Corbetta and Shulman 2002; Connor et al. 2004) were extended by the assumption of a fundamental value-driven attention mechanism (Anderson 2013; for a recent review, see Failing and Theeuwes 2017). This mechanism is discernible not only in stimuli inherently carrying salience, but also in stimuli associated with motivational valence, all sharing similar attentional prioritization. In line with this account, not only physical stimulus features but also emotional and motivational factors have been demonstrated to determine increased salience of certain stimuli and directly impact attention and visual processing capacities (e.g., Zeelenberg et al. 2006), resulting in a facilitated sensory encoding at initial processing stages (e.g., Della Libera and Chelazzi 2006). Stimuli of particularly high inherent salience are faces, for which involuntarily capture of attention and preferential processing has been documented, presumably due to their crucial role in human social interactions. This face-superiority effect has been reliably demonstrated on a behavioral level in object recognition/perception tasks (e.g., Langton et al. 2008), and moreover in studies employing visual search tasks or attentional blink paradigms including facial expressions of emotions (Eastwood et al. 2001; Anderson 2005; for a review, see Vuilleumier 2005; Calvo and Lundqvist 2008). Particularly, facial expressions of emotions convey various types of relevant information in social interactions (for a review, see Frith 2009) and are considered as evolutionary prepared stimuli (e.g., Vuilleumier and Pourtois 2007). Faces with and without emotional expressions are thus ideal stimuli in experiments investigating inherent versus associated salience effects as they allow for a direct comparison within an overall relevant stimulus domain.

Due to their high temporal resolution, event-related brain potentials (ERPs) allow segregating different processing stages and therefore gaining insights to the mechanism underlying the face superiority effect as well as the processing advantage of facial expressions of emotions over time. By means of ERPs, several studies indicated that the processing of facial expressions of emotion elicit amplified neural responses compared to other visual stimuli such as pictures of affective scenes or written words of emotional content (Schacht and Sommer 2009; Bayer and Schacht 2014). Attentional priority for facial expressions of emotion and their sustained preferential processing over neutral faces is reflected in several dissociable ERP components (e.g., Schupp et al. 2004; Rellecke et al. 2012). Especially two ERP components have been linked to subsequent stages of emotion processing in humans: the EPN and the LPC. The Early Posterior Negativity (EPN), a relative negativity over posterior electrode sites, typically starting around 150-200 ms after stimulus onset (e.g., Junghöfer et al. 2001; Rellecke et al. 2011), has been suggested to reflect enhanced sensory encoding of facial expressions of emotion. The EPN is typically followed by the Late Positive Complex (LPC) or Late Positive Potential (LPP, e.g., Cuthbert et al. 2000; Schupp et al. 2004) over centro-parietal electrodes, starting around 300 ms after stimulus onset (e.g., Rellecke et al. 2011). This long-lasting ERP response has been assumed to reflect higher-order elaborate and evaluative processes (for a review, see Olofsson et al. 2008; Schacht and Sommer 2009; Rellecke et al. 2011). In addition, two earlier components were recently found to be modulated by emotional expressions. First, the P1 component, is peaking around 100 ms after stimulus onset, consisting of bilateral occipital positivities and reflecting the activation of extrastriate visual areas via selective attention (Di Russo et al. 2003). Enhanced P1 amplitudes have been reported for emotional facial expressions in comparison to neutral facial expressions (e.g., Batty and Taylor 2003; Rellecke et al. 2011), indicating that emotional salience impacts early perceptual encoding. Second, the N170 has been functionally linked to holistic face perception, consisting in a negativity over temporo-occipital electrodes (e.g., Bentin et al. 1996). As the evidence of N170 modulations by emotional expressions is inconsistent (for reviews, see Rellecke et al. 2013; Hinojosa et al. 2015), the question whether configural and emotional features of a face are processed independently (Bruce and Young 1986) could yet not finally be answered.

Facial expressions of emotions, as well as other stimuli of emotional content, carry an increased motivational salience, e.g., angry faces trigger the avoidance system, while happy faces might carry reward in social interactions. Previous studies have demonstrated that even neutral faces gain salience through associated emotional context information (Suess et al. 2013; Wieser et al. 2014). However, in particular motivational salience might arise from a variety of other sources, driven by first, an explicit motivational context or second, by acquired associations. Contexts might determine motivational dispositions – e.g., the readiness to act in given situations – as they can confront a person with appealing opportunities and daunting obstacles (Scheuthle et al. 2005) and thus directly influence behavior. An increase of the motivational salience of a given context can be generated by introducing reinforcements as incentives (Meadows et al. 2016). In a recent ERP study, Wei and colleagues (Wei et al. 2016) showed that the expectation of monetary gain – indicated by motivationally relevant cues – impacted stimulus processing over consecutive stages from sensory encoding (*EPN*) to higher-order evaluation (*P3*/*LPC*). Interestingly, motivational incentives have been recently demonstrated to affect stimulus processing even before effects of spatial attention (Bayer et al. 2017). In addition, a “*cue-P3*” component directly elicited after cue onset with enhanced amplitudes for reward-indicating as compared to loss-indicating cues was reported (Zheng et al. 2017).

Driven by the compelling evidence for impacts of motivational contexts and inherent emotional valence, the question arises under which conditions salience can be acquired. The value-driven attention mechanism proposed by Anderson (Anderson 2013) incorporated this question suggesting that processing advantages are not restricted to stimuli of emotional content (e.g., facial expressions of emotion), but also hold for stimuli that have been associated with reward, even if these are inherently nonsalient or task-irrelevant. A fruitful approach to test this assumption is provided by associative learning paradigms that allow the investigation of the influences of acquired salience without interference with stimulus-driven salience. Aiming at a direct comparison between inherent and associated saliences, Hammerschmidt and colleagues (Hammerschmidt et al. 2017) reported that explicit reward-associations to inherently neutral faces elicited increased P1 responses during delayed testing. The elicitation of typical emotion-related ERP components at longer latencies (EPN and LPC), was, however, restricted to facial expressions of emotion. In contrast, employing a highly similar learning paradigm as in the study by Hammerschmidt et al. (Hammerschmidt et al. 2017), Rossi and colleagues (Rossi et al. 2017) detected an increase of the P3 to reward-associated unknown single letters from unfamiliar alphabets. Importantly, the processing advantage reported for stimuli associated with motivational salience is not restricted to rewards but has also been demonstrated for associations with aversive events (Stolarova et al. 2006; Hintze et al. 2014) or monetary loss (Rossi et al. 2017), mainly present on the perceptual level.

ERPs reflect processing differences on the neural level but cannot directly be linked to physiological arousal – one of the key components of emotions (Scherer 2005; Scherer 2009; Lang and Bradley 2010). Physiological arousal is reflected amongst other indicators in changes of the pupil size, which have been related to norepinephrine release in the locus coeruleus (Berridge and Waterhouse 2003; Einhäuser et al. 2008; Gilzenrat et al. 2010; Laeng et al. 2012; Murphy et al. 2014). Therefore, pupil activity can be used as a measure of attentional, cognitive and emotional processing (Smallwood et al. 2011; Kang et al. 2014), with increased pupil size in response to emotionally arousing pictures (Bradley et al. 2008) and auditory stimuli (Partala and Surakka 2003). In particular, inherently angry faces paired with an angry body induced larger pupil dilations than fearful and happy face-body pairs (Kret et al. 2013). Moreover, motivational modulations through outcome associations, in addition to stimuli of inherent emotional salience, can also increase pupil size, demonstrated for both reward (e.g., Massar et al. 2016) and loss incentives (Pulcu and Browning 2017). Interestingly, modulations of pupil dilation further depend on task difficulty, manipulated through mental effort (Mathôt et al. 2015; Peysakhovich et al. 2015), and decision uncertainty (Kahneman 1973; Satterthwaite et al. 2007; Brunyé and Gardony 2017; Urai et al. 2017), with greater pupil dilations occurring with increasing task difficulty. The parallel measurement of ERPs, pupil dilations and behavioral data might help elucidate the multiple components involved in emotion processing (e.g., Grandjean et al. 2008).

In line with Anderson’s assumption (Anderson 2013) of a value-driven attention mechanism, suggesting shared mechanisms of inherent bottom-up stimulus attention and context- or learning-based salience effects, previous research clearly indicated that both emotional and motivational aspects have a direct impact on visual stimulus processing. Nevertheless, the specific conditions, under which learning mechanisms or different contexts can modify a certain stimulus’ salience, are not fully understood, presumably contributing to heterogeneous findings in the past.! Despite the great progress in this area of research, there are a number of outstanding open questions that have not sufficiently been addressed: Firstly, effects of associated motivational salience occurred during several processing stages mainly in explicit associative learning paradigms (e.g., Stolarova et al. 2006; Hintze et al. 2014; Hammerschmidt et al. 2017; Rossi et al. 2017). However, it seems reasonable that motivation or emotion-based salience might have been acquired implicitly, that is without explicit knowledge about the hedonic value of the certain stimulus. Hence, one of the yet unresolved questions is whether implicit and explicit associations of motivational salience have similar effects on stimulus processing. Implicit learning is generally linked to participants/learners’ problems with an explicit recall (Berry and Dienes 1993), often characterized as a ‘complex form of priming’ (Cleeremans et al. 1998). Further, it was argued that implicit representations possibly need more time and cognitive resources to be generated than information learned explicitly (Batterink and Neville 2011). Recently, it could be demonstrated that reward associations have a direct impact on spatial attention – even when presented implicitly (Bourgeois et al. 2016). Secondly, it remained open whether the impacts of associated gain and loss might be symmetric under conditions of equalized outcomes, as successful learning usually implies an increase of gain in parallel to reduced losses (e.g., Hammerschmidt et al. 2017; Rossi et al. 2017).

The main aim of the current study was to investigate potential effects of implicitly learned associations of motivational salience to neutral facial stimuli in direct comparison to effects elicited by inherent facial expressions of emotion. To this aim, we employed a prime-face matching task with subliminal prime presentation, implementing performance at chance level and thus an equalization of performance-dependent gain, loss, or zero-outcome conditions. During the learning session, colored cues were presented at the beginning of each trial, indicating the motivational condition which was kept constant for each of the inherently neutral target faces. During the test phase, the same task was employed, however without any performance-depended monetary incentives and feedback. In addition to the previously learned neutral faces, facial expressions of emotion of novel identities were presented, allowing for a comparison of effects driven by associated motivational and inherent emotional salience. We collected ERP and pupil size data during the learning and test sessions with the main aim to test the impact of motivational contexts on subsequent stimulus processing (cf., Wei et al. 2016) and to allow the investigation of the temporal characteristics and autonomous physiological correlates of association-related effects on the following day. We expected that the cue-indicated reward or loss would boost sensory processing of task-relevant face stimuli in the visual cortex (Bayer et al. 2017), resulting in enhanced P1 amplitudes after target face onset. Aiming at expanding the findings by Zheng and colleagues (Zheng et al. 2017) that showed augmented P3 amplitudes elicited by reward-indicating visual cues, we further tested potential modulations of cue-evoked ERP potentials by different motivational contexts. As the incentive values of the cue stimuli were made explicit to our participants, these simple symbolic stimuli might carry increased salience as stimuli of emotional/motivational content and thus trigger increased amplitudes of EPN and LPC components. Pupil dilations should be increased in condition of high motivational salience (Massar et al. 2016; Pulcu and Browning 2017). For faces associated with monetary gains on the previous day, we expected increased amplitudes of early ERP components (e.g., P1; Hammerschmidt et al. 2017). Loss-associations might trigger similar effects as gain-associations as both incentive conditions were equalized – in terms of frequency of occurrence and amount of monetary outcome – during the learning session. Faces with happy and particularly with angry expressions should elicit larger EPN and LPC amplitudes than neutral expressions (e.g., Schupp et al. 2004; Schacht and Sommer 2009; Rellecke et al. 2011. For pupil dilations, we expect an increase for angry compared to happy and neutral expressions (Kret et al. 2013). Pupil dilations to neutral faces associated with motivational salience the day before might show no increase due to the absence of arousing motivational context.

## Materials and Methods

### Participants

Data was collected from fifty-five participants. Seven participants were excluded due to EEG artifacts in either the learning or test phase, and four due to strategies that successfully countered visual masking during the face-matching task (the performance exclusion criterion was defined as an individual performance-dependent bonus exceeding average bonus ±2SDs across participants in the learning session). The remaining forty-four participants (21 female) were ranging in age between 18 and 32 years (mean age = 24.0 years, *SD* = 3.5), with normal or corrected-to-normal vision and without neurological or psychiatric disorders according to self-report. Forty-two participants were right-handed (according to Oldfield 1971). Participants received 8 euro per hour or course credit; in addition, the individual monetary bonus achieved during the learning phase was disbursed.

### Stimuli

Facial stimuli were selected from the Karolinska Directed Emotional Faces (KDEF) database (Lundqvist et al. 1998). Twelve colored pictures of faces (6 female, 6 male) with neutral facial expressions were used as target faces. The same pictures served as primes in matching trials; additional pictures of neutral faces (6 female, 6 male) were used as nonmatching primes. An ellipsoid mask surrounded all facial stimuli within an area of 130 × 200 pixels (4.59 × 7.06 cm, 4.6 × 7.1°) in order to eliminate hair, ears and clothing and leave only the face area visible.

For the learning phase, diamond-shaped cues of 120 × 120 pixels (3.18 × 3.18 cm) were generated that indicated the outcome category (reward, loss, zero outcome) of the given trial in three different equi-luminant colors (blue, pink, and brown). Grey circles were used as feedback stimuli (248 × 248 pixels, 5 × 5 cm) indicating the amount of monetary outcome won or lost in the preceding trial in the corresponding cue color.

For the test phase, twelve novel identities with facial expressions of emotion (happy, neutral, angry, *N* = 36 colored pictures) were presented in addition to the neutral faces which were presented during the learning phase the day before both as target faces and matching primes. Another twelve new identities (6 female, 6 male) showing facial expressions of emotion (happy, neutral, angry, *N* = 36 colored pictures) were used as prime stimuli in non-matching trials.

For each face stimulus (in total *N* = 96), a scrambled version was generated and used as mask for the preceding primes. All facial stimuli were matched offline for luminance (according to Adobe Photoshop CS6™), *F*(23,72) = 0.873, *p* = 0.631. All stimuli were presented in the center of the screen on a light gray background.

### Procedure

The study was conducted in accordance with the Declaration of Helsinki and approved by the local ethics committee of the Institute of Psychology at the University of Göttingen. Participants were informed about the procedure of the study and gave written informed consent prior to both phases of the experiment. The study consisted of a learning and a test phase, which were completed on two subsequent days. Participants were seated in a dimly lit, sound-attenuated room, in front of a computer screen (refresh rate 100 Hz) at a distance of 57 cm. Participants placed their chin and forehead on a head rest in order to avoid movements and ensure correct recording of pupil sizes. After pupil diameter calibration, participants received detailed instructions about the experimental task.

In the learning phase, twelve inherently neutral faces were implicitly associated with monetary gain, loss, or no outcome via an associative learning paradigm. At the beginning of each trial, a diamond-shaped cue indicated the monetary outcome context condition (gain, loss, or neutral: no gain/loss). The assignment of the cue’s color was fixed for each participant but counterbalanced across participants. The meaning of the cues and the feedback scheme was explained prior to the experiment. Participants were asked to decide whether the identity of the presented target face was matching the preceding prime face – irrespective of the presented cue. In the gain condition, the correct classification of the face-matching task was awarded with +50 cents (incorrect classifications = 0 cents). A correct classification in the loss condition prevented the participants from the loss of money (0 cents), whereas an incorrect classification led to a loss of 50 cents. For the neutral condition, feedback was either +0 cents (correct classification) or -0 cents (incorrect classification). Responses were given by a button press; correct/incorrect-buttons as well as prime-target assignments were counterbalanced, but consistent within one participant. In the face-matching task, prime and target faces differed in 50% of the trials in identity, but were always matched with respect to gender. In case the participant missed to answer a trial within 5000 ms, 70 cents were removed from the bonus. Stimuli were presented blockwise with a total of 20 blocks. Each block consisted of the 12 target faces with neutral expressions presented twice in randomized order, paired with a matching (50%) or a non-matching (50%) prime, resulting in 480 trials in total. Importantly, the cue-target face associations remained stable during the learning phase for each participant, but were counterbalanced in order to exclude any potential effects of physical stimulus features on the ERP components of interest. At the beginning of each trial (see Figure 1), a fixation cross was presented in the center of the screen for 1000 ms, followed by the diamond-shaped cue, which was visible for 500 ms. Subsequently, a fixation cross was shown for 200 ms followed by the prime face for 10 ms. The mask appeared for 200 ms followed by a fixation cross for 200 ms. The target face was shown up to 5000 ms, disappearing with button press. The feedback was displayed for 1000 ms. Blocks were separated by a self-determined break, in which the current amount of the individual bonus was displayed. Participants started with a base pay of 10 euro and achieved an individual monetary bonus according to their performance ranging between -11 and 18 euro (*mean* = 1.11 euro, *SD* = 5.98 euro); participants finishing the learning session with a negative balance received the full base payment of 10 euro.

**Figure 1.**
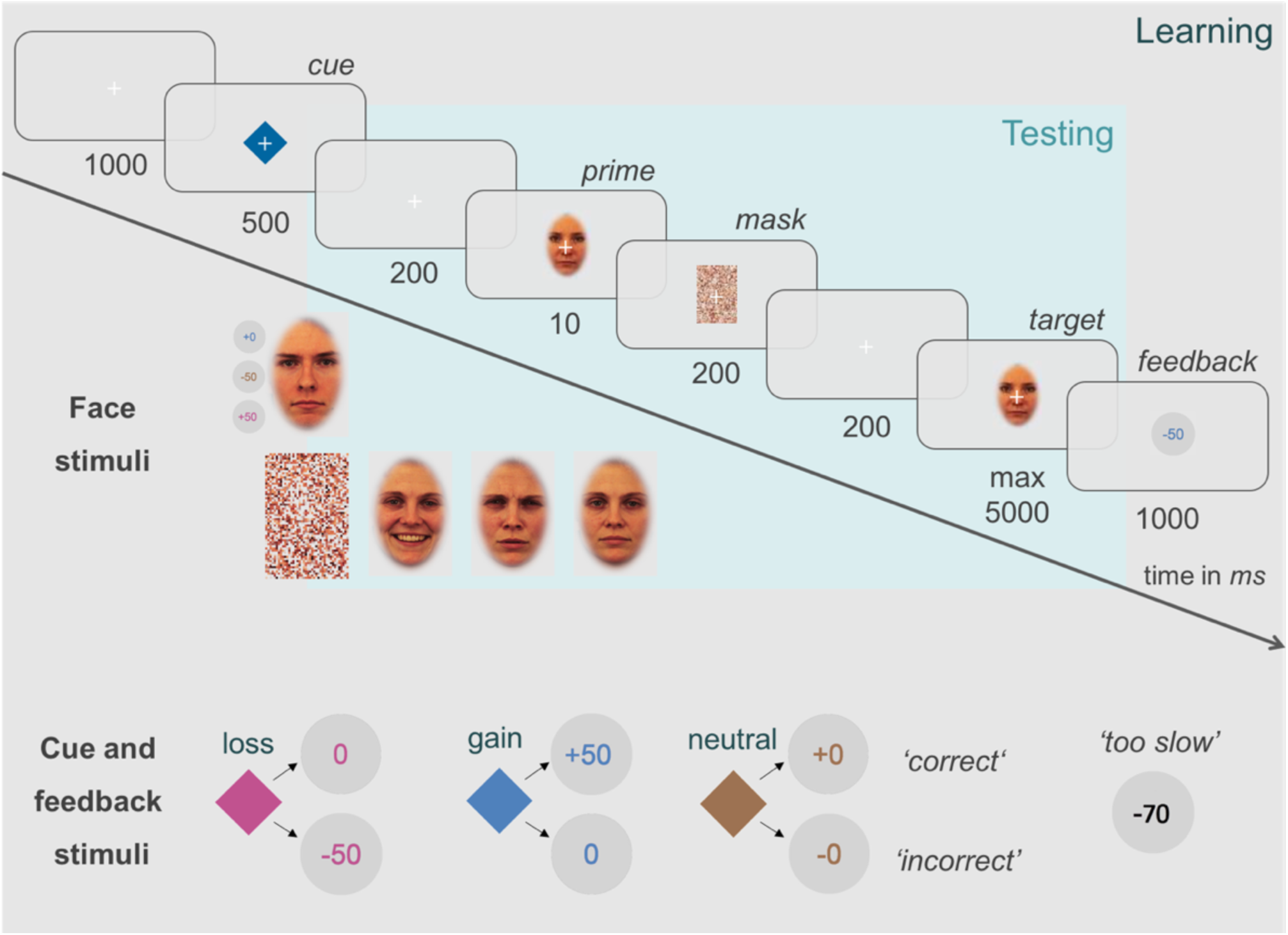
Trial scheme of the learning and test session with detailed time sequence.

In order to check whether the associations of the presented cue and the target face remained implicit, a manipulation check was implemented at the end of the learning phase. The twelve target face identities were presented simultaneously, randomly arranged on the computer screen. The participants were asked to explicitly assign them to one of the three outcome contexts (gain/neutral/loss). This task was repeated about 30 minutes later.

The test phase took place on the following day, to allow for memory consolidation. The face-matching task remained constant, however, no cue or corresponding feedback was provided, and participants could not win or lose any money. The test phase consisted of two different types of facial stimuli presented blockwise. Half of the blocks consisted of the twelve neutral target faces, which were implicitly associated with monetary outcome context the day before. The other half of the blocks consisted of twelve novel identities with emotional facial expressions (4 for, happy, neutral, and angry, respectively) serving as target face and primes in matching trials, and twelve additional novel identities with emotional expressions (4 for, happy, neutral, and angry, respectively) serving as primes in the non-matching trials. Target and prime faces always matched with respect to gender and emotional expressions. As in the learning phase, each target face was presented twice with a matching and a nonmatching prime in randomized order (N = 48 blocks). The trial scheme was identical to the learning session, except that cues and feedback stimuli were excluded (see Figure 1). Each block was repeated ten times in randomized order, resulting in 20 blocks and 960 trials in total per face condition. The blocks were separated by breaks of self-determined length. Again, a manipulation check was conducted at the end of the test phase: all 24 target face identities with neutral expressions (from both blocks with previously learned and inherent facial expressions) were presented on the computer screen in random order. The participants were asked for each face whether it was presented during the learning phase the day before or during the test phase for the first time.

### Acquisition and pre-processing of ERP and pupil data

The EEG was recorded from 64 electrodes, placed in an electrode cap (Easy-Cap, Biosemi, Amsterdam, Netherlands) according to the international 10-20 system (Pivik et al. 1993). The common mode sense (CMS) electrode and the driven right leg (DRL) passive electrodes were used as reference and ground electrodes (http://www.biosemi.com/faq/cms&drl.htm). Six external electrodes were used: Two on the left and right mastoids respectively, and four external electrodes were placed on the outer canthi and below the eyes to record eye movements and blinks. Signals were recorded at a sampling rate of 512 Hz and a bandwidth of 102.4 Hz (http://www.biosemi.com/faq/adjustfilter.htm), offline filtered with a Low Cutoff (0.03183099 Hz, Time constant 5 s, 12 dB/oct), a High Cutoff (40 Hz, 48 dB/oct) and a Notch Filter (50 Hz). Data was processed using BrainVision Analyzer (Brain Products GmbH, Munich, Germany). Data was down-sampled to 500 Hz, average-referenced and corrected for ocular artifacts (blinks) using Surrogate Multiple Source Eye Correction with default parameters (MSEC; Ille et al. 2002) as implemented in BESA (Brain Electric Source Analysis, MEGIS Software GmbH, Gräfelfing, Germany). Application of Surrogate MSEC is detailed in Scherg (Scherg 2003). The continuous EEG signal of the learning phase was segmented into epochs of 2310 ms, starting 200 ms before cue onset and referred to a 200 ms pre-cue baseline. The continuous EEG signal of the test phase was segmented into epochs of 1610 ms, starting 200 ms before prime onset and referred to a 200 ms pre-prime baseline. Based on previous research (Hammerschmidt et al. 2017), time windows and regions of interest (ROIs) electrodes for ERP components were chosen as follows for the learning session (related to cue onset): P1 cue: 75-125 ms; EPN cue: 200-300 ms; LPC cue: 350-500 ms; P1 fixation cross1: 585-635 ms; P1 prime/mask: 760-810 ms; P2 prime/mask: 885-935 ms; P1 fixation cross2: 985-1035ms; P1 target: 1185-1235 ms; N170 target: 1240-1290 ms; EPN target: 1310-1460 ms; LPC target: 1460-1810 ms. For the test session (related to target face onset): P1: 75-125 ms, N170: 130-180 ms, EPN: 200-350 ms, P3: 200-350 ms, LPC: 350-700 ms. ERPs were quantified as most positive peak using peak detection (P1 at O1 and O2, reference electrode: O2; N170 at P9 and P10, reference electrode: P10; P2, O1 and O2, reference electrode: O2) or mean amplitudes (EPN at P9, P10, Iz, Oz, O1, O2, PO7, and PO8; LPC at Pz, P1, P2, CPz, and POz).

Pupil diameter was recorded binocularly using a desktop-mounted eyetracker (EyeLink 1000, SR Research) at a 500 Hz sampling rate. Prior to the experiment, pupil diameter was calibrated with an artificial pupil placed on the lid of the left eye of the participants to set the baseline for the measurement of the pupil dilation size. Offline, analyses of pupil diameter were performed using Matlab. Trigger codes of pupil and EEG data were synchronized.

Data from two subjects were excluded due to technical failure of the eye tracker in the learning or test phase, respectively. For each participant and the learning and test sessions separately, artifacts were identified as samples in which the difference in pupil size to the subsequent sample was higher than 0.1 mm or the difference in pupil size from the median across the session was higher than 1 mm. Artifacts were interpolated. Eleven subjects had to be excluded after artifact correction due to excessive artifacts that could not be interpolated in either the learning or the test session. The remaining pupil size data was segmented into epochs from 200 ms prior to cue (learning session/prime (test session) onset to 7000 ms after. For each subject and condition, pupil size time courses were averaged across both eyes and correct and incorrect responses, and corrected to a baseline 200 ms before cue (learning session)/prime (test session) onset. Mean pupil size between 1500 and 4000 ms after cue/prime onset (based on the response latency after cue onset measured by Bayer et al. 2017) was computed for each subject and condition. One additional subject was excluded because the measured pupil size exceeded the average across subjects by more than 10 SD.

### Data analyses

All parameters – reaction times (RTs), accuracy (in percent), ERP peaks or mean amplitudes, and pupil diameter – were analyzed with repeated-measures (rm) ANOVAs, separately for the learning session and test session. Outliers were identified as reaction times (RTs) below 200 ms or exceeding +2SDs from the mean per condition and were excluded from behavioral data analysis. RmANOVAs on data from the learning session included the factor Motivation (gain, neutral, and loss). Data from the test phase were analyzed in separate rmANOVAs, including the factor Motivation (gain, neutral, and loss) for learned faces or the factor Emotion (happy, neutral, and angry) for novel faces with emotional expressions. Accuracy deviations from chance level, across the sample and on the individual subject level, were analyzed using the exact test for equality of several binomial proportions to a specified standard (Krishnamoorthy et al. 2004; Unakafov, 2017). All post-hoc pair-wise comparisons were Bon-ferroni-corrected.

## Results

### Effects of Motivational Context in the Learning Phase

#### Behavioral Data

Descriptive values for behavioral performance measures of the learning session are provided in Table 1. Accuracy on the face-matching task during the learning phase was at 50% chance level (not different from the expected random binomial distribution with 0.5 probability, *p* > 0.05, Bonferroni-corrected), and was not impacted by the factor Motivation, *F*(2,86) = 0.149, *p* = 0.850, *η^2^_p_* = 0.003. Mean reactions times (RTs) of the learning phase significantly differed as a function of the factor Motivation, F(2,86) = 24.929, *p* < 0.001, *η^2^_p_* = 0.367, with increasing RTs from neutral to gain- and loss-context, and loss to gain context trials, all Fs(1,43) > 11.206, all *ps* < 0.006, all *η^2^_p_* > 0.207.

**Table 1.** Mean reaction times in ms, accuracy in task and manipulation check in %, during/after face-matching task in the learning session (SEMs in parentheses), contrasted for factor levels of Motivation.

|  | Learning Session |  |  |  |
| --- | --- | --- | --- | --- |
|  | Face-Matching Task |  | Manipulation Checks |  |
|  | RTs | Accuracy | 1<sup>st</sup> Check | 2<sup>nd</sup> Check |
| Gain | 1019 (49) | 51 (0.7) | 57 (3.3) | 55 (3.9) |
| Neutral | 960 (44) | 51 (0.6) | 48 (4.6) | 48 (3.9) |
| Loss | 1079 (51) | 51 (0.7) | 45 (3.4) | 47 (3.7) |

Correct assignments of the target faces to motivation conditions – obtained directly after the learning phase (1^st^ check) and after 30 minutes delay (2^nd^ check) – were above 33% chance level for gain-and neural-associated faces (p < 0.05, Bonferroni-corrected, the exact test for equality of several binomial proportions to a specified standard), but did not reach significance for loss-associated faces, without any performance improvement after 30 minutes delay, *F* < 1.

#### ERP Data

*ERPs elicited by motivational cues.* EPN mean amplitudes between 200 and 300 ms after cue onset differed as a function of Motivation, F(2,86) = 7.960, *p* = 0.001, *η^2^_p_* = 0.156, for gain-compared to neutral-, F(1,43) = 10.295, *p* = 0.009, *η^2^_p_* = 0.193, and loss-compared to neutral-related trials, F(1,43) = 14.837, *p* < 0.001, *η^2^_p_* = 0.257. LPC mean amplitudes between 350 and 500 ms after cue onset were also modulated by Motivation, F(2,86) = 37.755, *p* < 0.001, *η^2^_p_* = 0.468, with enhanced amplitudes for gain-compared to neutral-, F(1,43) = 52.145, *p* < 0.001, *η^2^_p_* = 0.548, for loss-compared to neutral-, F(1,43) = 26.100, *p* < 0.001, *η^2^_p_* = 0.378, and for gain-compared to loss-related trials, F(1,43) = 22.067, *p* < 0.001, *η^2^_p_* = 0.339. The P1 elicited by motivational cues was not impacted by the factor Motivation (see Figure 2).

As can be seen in Figure 2, the impacts of motivational incentives were long-lasting. Therefore, ERPs between cue and target face presentation were analyzed to investigate potential impacts of motivational context. The P1 component following the first fixation cross after cue presentation was modulated by the Factor Motivation, F(2,86) = 8.752, *p* = 0.001, *η^2^_p_* = 0.169, with enlarged peak amplitudes for reward-compared to neutral-, F(1,43) = 16.513, *p* < 0.001, *η^2^_p_* = 0.277, and loss-compared to neutral-related trials, F(1,43) = 7.115, *p* = 0.033, *η^2^_p_* = 0.142. Motivation further influenced the P1 component following prime/mask, F(2,86) = 13.959, *p* < 0.001, *η^2^_p_* = 0.245, with larger positivities for reward-compared to neutral-, F(1,43) = 25.947, *p* < 0.001, *η^2^_p_* = 0.376, and loss-compared to neutral-related trials, F(1,43) = 10.699, *p* = 0.006, *η^2^_p_* = 0.199. The visual P2 following prime/mask was also modulated by the Factor Motivation, *F*(2,86) = 5.934, *p* = 0.005, *η^2^_p_* = 0.121, with enhanced peak amplitudes for loss-compared to neutral-related trials, F(1,43) = 10.981, *p* = 0.006, *η^2^_p_* = 0.203. The fixation cross response following the prime/mask was not modulated by the factor Motivation anymore (see Figure 2, panels A and B).

**Figure 2.**
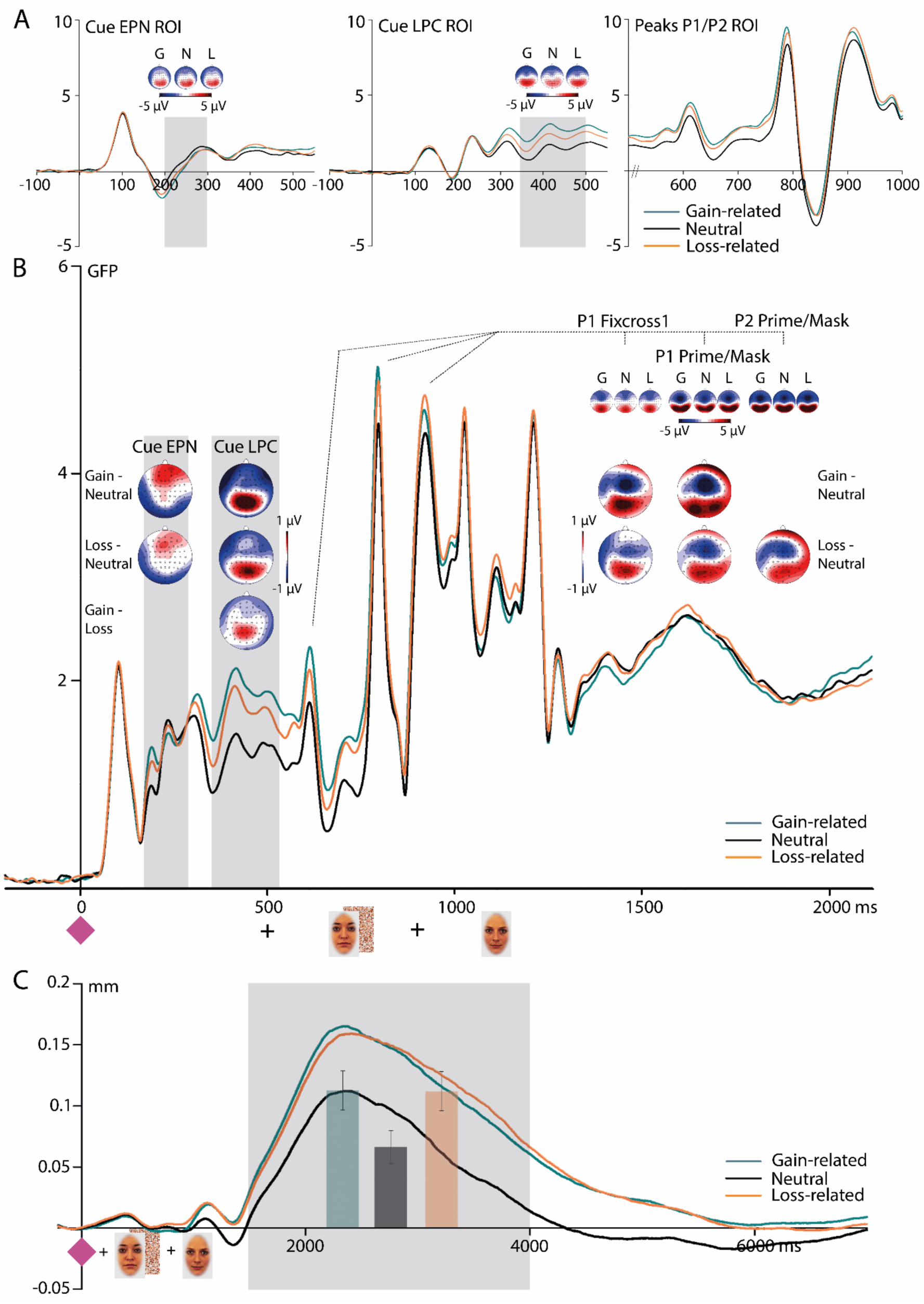
*ERP effects of the learning phase for Cue-EPN and Cue-LPC for associated faces and the following peaks. **A**: Regions of interest (ROIs) for the corresponding analyses. **B**: GFP wave form of a complete trial for reward-, neutral- and loss-related faces including ERP topography of raw distributions (small topographies) and differences between indicated motivation categories. Highlighted areas display the time windows of Cue- ERP analyses, P1/P2 peaks of the after-cue/pre-target face interval were analyzed with peak detection. **C**: Pupil dilation responses to gain-, neutral-, and loss-related contexts, the highlighted area displays the time window of pupil dilation analysis with means and SEMs embedded as bar chart*.

*ERPs to Target Faces.* According to rmANOVAs, modulations of peak amplitudes for P1 and N170 components and mean amplitudes for EPN and LPC components by implicitly associated motivational salience were absent. *Pupil dilations.* For pupil dilation data of the learning phase, an rmANOVAs showed a significant within-subjects effect of Motivation, F(2,58) = 32.871, *p* < 0.001, *η^2^_p_* = 0.531, with increased pupil diameters for gain-compared to neutral-, F(1,29) = 43.413, *p* < 0.001, *η^2^_p_* = 0.6, and loss-compared to neutral-related trials, F(1,29) = 33.466, *p* < 0.001, *η^2^_p_* = 0.536 (see Figure 2, panel C).

### Effects of Associated Motivational and Inherent Emotional Salience in the Test Phase

#### Behavioral Data

Descriptive values for behavioral performance measures of the test session are provided in Table 2. In contrast to the learning session, the accuracy on the face-matching task in the test phase across the sample of 44 subjects was slightly above the 50% chance level (*Ms* = 51-53%, *p* < 0.05, Bonferroni-corrected). In particular, five subjects showed a significant accuracy above (4 subjects, accuracy 58-65%) or below chance level (one subject, 40%) across all three motivational conditions, for the previously associated faces (*p* < 0.05). Similarly, six subjects (four same as for the motivational conditions) showed above chance accuracy for novel faces across all three emotional conditions (58-70%). Accuracy was not impacted by the factors Motivation/Emotion, and did not differ between conditions (learned faces /novel faces), *Fs* < 1.4. During the test phase, RTs were not modulated by the Factor Motiva-tion/Emotion, *Fs* < 1.

**Table 2.** Mean reaction times in ms, accuracy in task and manipulation check in %, during/after face- matching task in the test session (SEMs in parentheses), contrasted for all factor levels of Motivation/Emotion. Man.-Ch. = Manupiulation Check.

|  | Test Session |  |  |
| --- | --- | --- | --- |
|  | Face Matching Task |  | Man. Ch. |
|  | RTs | Accuracy | Old/New |
| Reward | 986 (57) | 51 (1.0) | 87 (3.0) |
| Neutral | 985 (57) | 51 (0.8) | 80 (3.7) |
| Loss | 978 (56) | 52 (0.8) | 83 (3.7) |
| Happy | 1011 (58) | 53 (0.9) | 86 (3.2) |
| Neutral | 1006 (55) | 51 (0.9) | 78 (4.0) |
| Angry | 1014 (58) | 51 (1.0) | 89 (3.4) |

After the test phase, all 24 target faces from both learning and testing phase were presented to the participants (2 subjects did not complete the retrieval). They had to assign those to either the learned target faces from the day before or to the novel target faces with emotional expressions of the test phase (average performance: *M* = 84.0%, *SEM* = 2.5%) to control for familiarization with the target faces of the learning phase. The factor Motivation did not impact accuracy of learned target faces. For novel target faces with emotional expressions, a main effect of the factor Emotion was detected, F(2,82) = 4.173, *p* = 0.020, *η^2^_p_* = 0.092, with higher accuracy rates for angry compared to neutral expressions, F(1,41) = 7.280, *p* = 0.030, *η^2^_p_* = 0.151.

#### ERP Data

##### ERP Effects of Associated Motivational Salience

rmANOVAs on ERPs revealed a significant main effect of the factor Motivation on LPC mean amplitudes for inherently neutral faces associated with motivational salience, F(2,86) = 10.632, *p* < 0.001, *η^2^_p_* = 0.198, with increased amplitudes for gain-associated compared to neutral faces, F(1,43) = 18.792, *p* < 0.001, *η^2^_p_* = 0.304, and compared to loss-associated faces, F(1,43) = 8.880, *p* = 0.015, *η^2^_p_* = 0.171 (see Figure 3). P1, N170, and EPN amplitudes to associated faces were not influenced by the Factor Motivation, when tested in the a-priori defined time windows and ROIs.

##### Further ERP Effects of Associated Motivational Salience prior to the LPC component

The time window 200-350 ms after target face onset, which revealed no EPN modulation for associ- ated motivational salience, was visually reinspected (see Figure 3) as amplitude distributions and corresponding topographies bore a high resemblance to the LPC effect (350-700 ms) of associated motivational salience outlined above. Therefore, the time window was reanalyzed with the centro-parietal LPC ROI revealing effects of associated motivational salience, F(2,86) = 5.124, *p* = 0.008, *η^2^_p_* = 0.106, with enhanced amplitudes for gain-compared to neutral-associated faces, F(1,43) = 8.346, *p* = 0.018, *η^2^_p_* = 0.163.

**Figure 3.**
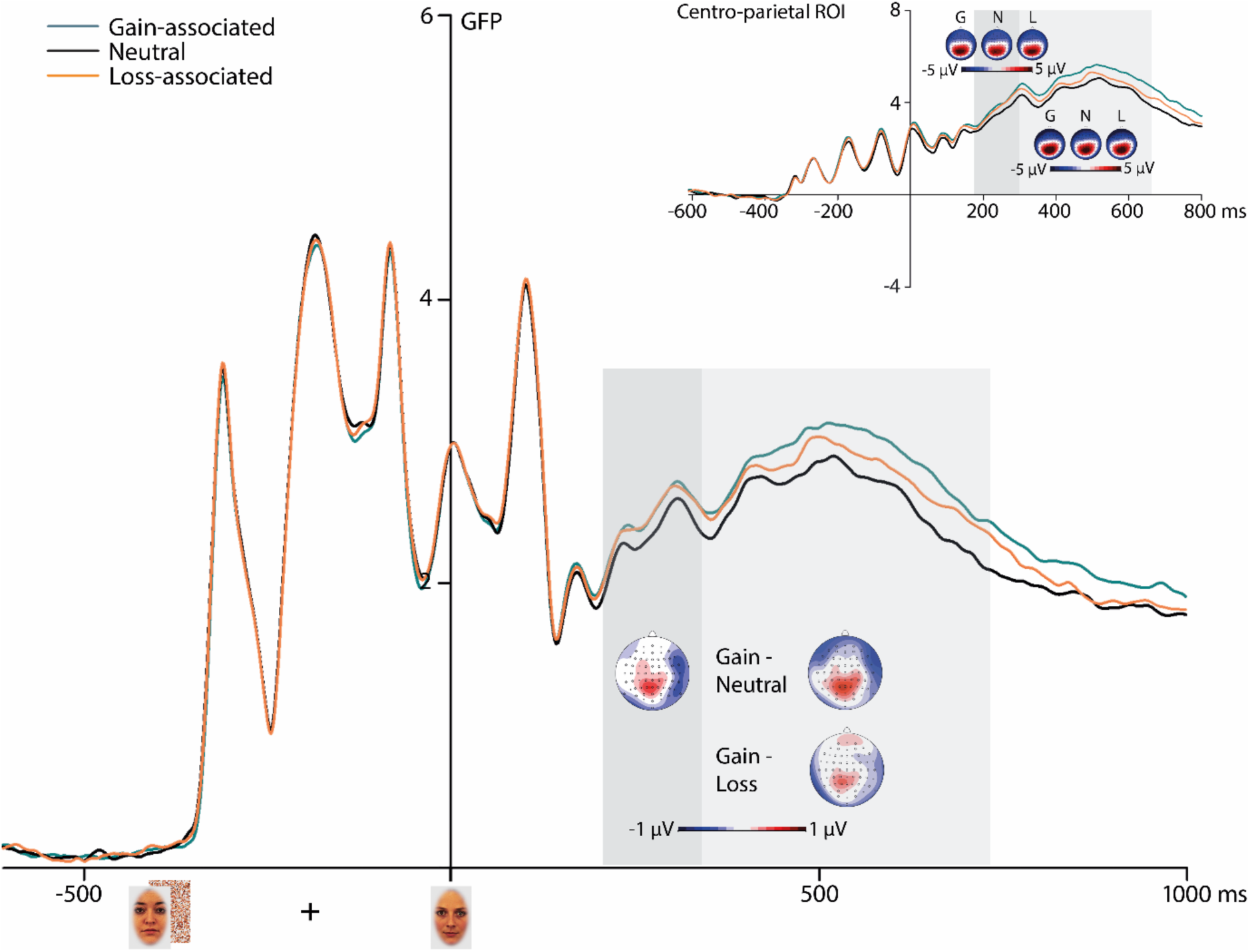
GFP wave form of a complete trial of the test session for gain-, neutral- and loss-associated faces including centro-parietal/LPC ROI and ERP topography of raw distributions (upper graph) and differences between indicated motivation categories. Highlighted areas display the time windows of analyses.

##### ERP Effects to Facial Expressions of Emotion in Novel Identities

N170 peak amplitudes to the target faces were significantly impacted by the factor Emotion, F(2,86) = 7.901, *p* = 0.001, *η^2^_p_* = 0.155, with enhanced negativities for angry compared to neutral, F(1,43) = 13.695, *p* = 0.003, *η^2^_p_* = 0.242, and happy expressions, F(1,43) = 8.941, *p* = 0.015, *η^2^_p_* = 0.172. EPN mean amplitudes of novel emotional expressions were significantly modulated by the Fac-tor Emotion, F(2,86) = 21.217, *p* < 0.001, *η^2^_p_* = 0.330, with enhanced amplitudes for happy compared to neutral, F(1,43) = 34.587, *p* < 0.001, *η^2^_p_* = 0.446, and for angry compared to neutral facial expressions, F(1,43) = 39.982, *p* < 0.001, *η^2^_p_* = 0.482. P1 peak and LPC mean amplitudes for novel faces with emotional expressions were unaffected by the Factor Emotion (see Figure 4).

**Figure 4.**
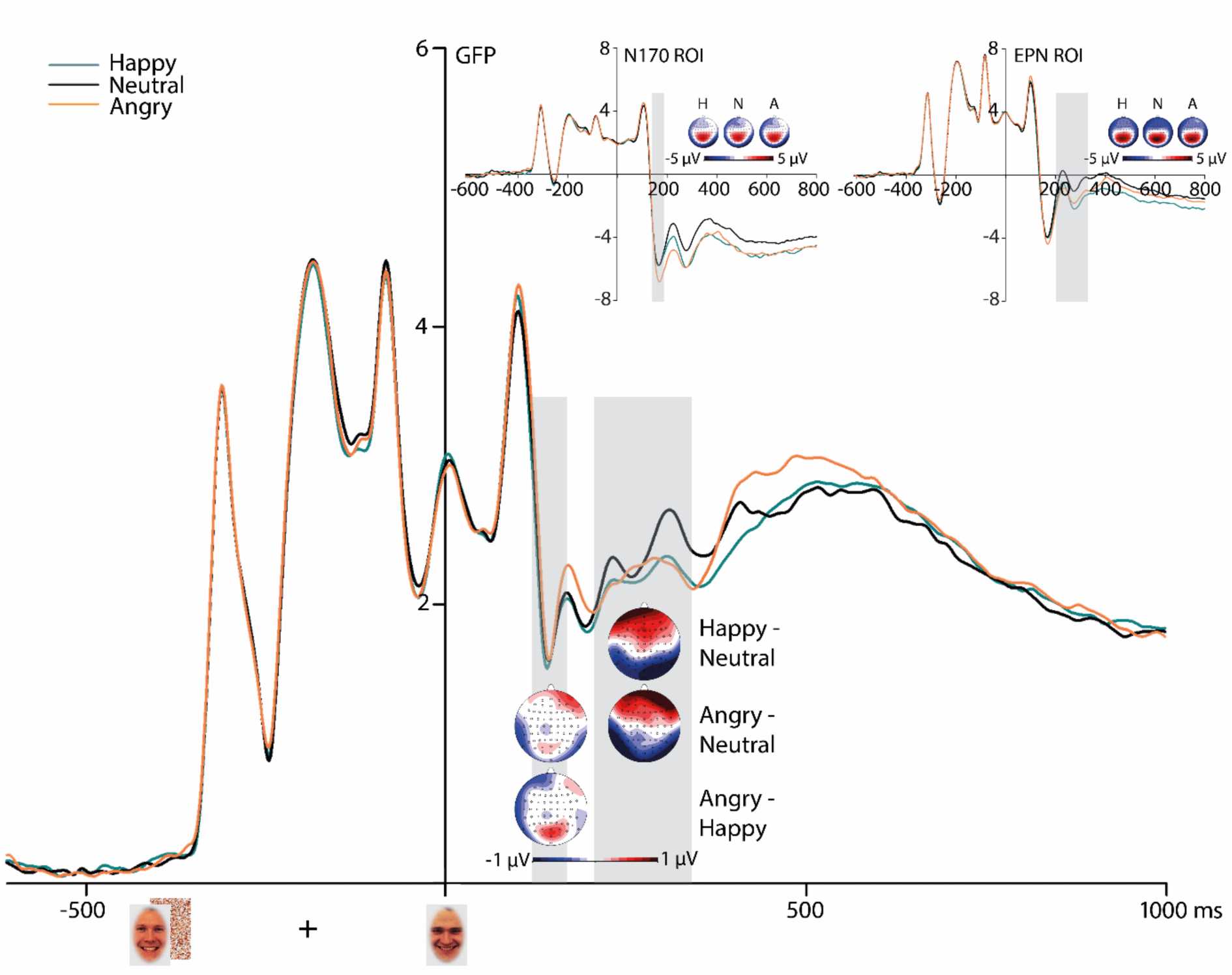
*GFP wave form of a complete trial of the test session for happy-, neutral- and angry faces including N170/EPN ROIs and ERP topography of raw distributions (upper graphs) and differences between indicated emotion categories. Highlighted areas display the time windows of analyses*.

##### Pupil dilations

An rmANOVA showed no significant within-subjects effect of associated motivational salience on pupil size, F(2,58) = 0.049, *p* = 0.950, *η^2^_p_* = 0.002. Pupil size in response to novel facial stimuli with emotional expressions did not significantly differ, according to an rmANOVA, F(2,58) = 0.705, *p* = 0.498, *η^2^_p_* = 0.024 (see Figure 5).

**Figure 5.**
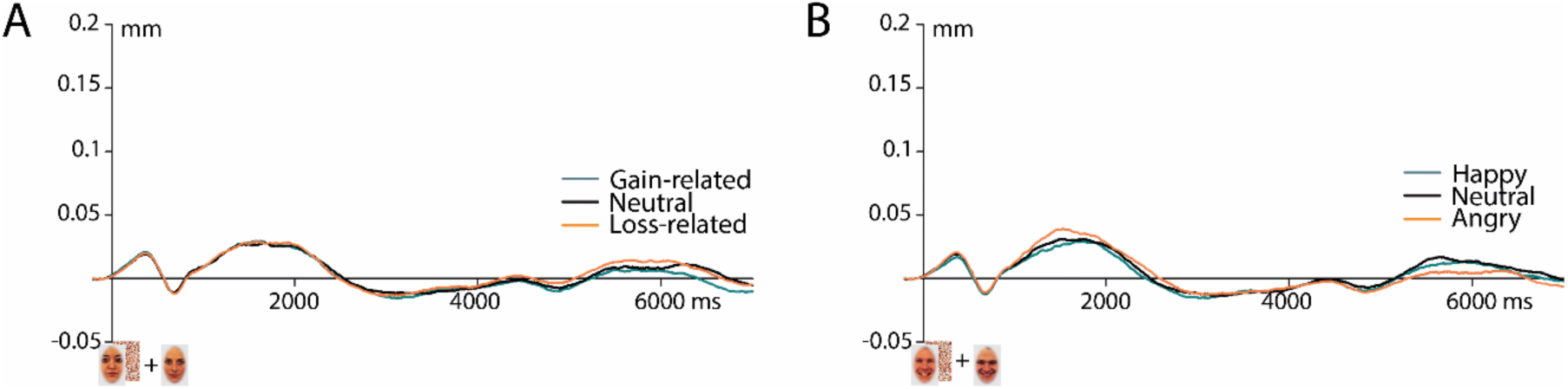
Pupil dilations during the test phase for A: previously associated and B: inherent emotional expressions.

##### Topography comparisons

As there is no previous evidence for emotion/motivation-related ERP modulations following a motivational cue, it is an exploratory question whether a P3 modulation or an EPN modulation could be expected prior to LPC modulations driven by the valence of the cue. To decide whether the ERP difference modulations between 200-300 ms after cue onset resemble an EPN distribution, topography comparisons were measured. To this end, the mean amplitude of all 64 electrodes was divided by global field power (GFP; Skrandies 1990) per condition respectively to extinguish amplitude differences. Difference of the particular conditions were measured and compared with the topography of an established ERP component via rmANOVAs with the factor Electrode (64) and the factor Topography (2). To compare the topography of the ERP modulation 200-300 after cue onset, the difference topography of gain minus neutral cues was compared with the difference topography of happy minus neutral expressions of the test phase. The topography x electrode interaction revealed no significant difference between these two topographies, *F* < 1. Similarly, the difference topography of loss minus neutral cues was compared to the difference topography of angry minus neutral expressions of the test phase. The topography x electrode interaction again failed significance between these two topographies, *F* < 1.325.

## Discussion

The main aim of the present study was to investigate whether implicitly learned associations of motivational salience result in a prioritized processing similar to what has been previously shown for explicit associations or inherent emotional salience (e.g., Hammerschmidt et al. 2017). Further, effects of motivational incentives on subsequent stimulus processing were examined during learning, while gain and loss were held equal in terms of their frequency and amount of monetary outcome. To address these aims, we implemented a multi-measure approach, considering ERPs as indicator of neural processing, pupil dilations as a correlate of arousal, and behavioral parameters as control variables. During a learning session, a sequential face-matching task using inherently neutral faces as subliminal and masked primes and supraliminal targets was employed, while motivational context was indicated by preceding cues and feedback about monetary outcome at the end of each trial. Importantly, target face assignments to motivational context were kept constant for each participant (but were counterbalanced between participants). On the following day, the previously associated faces were presented together with novel faces with expressions of emotion (happy, angry, and neutral faces) allowing for a direct comparison of potential effects driven by associated versus inherent salience during face processing.

### Implicitly acquired reward associations improve stimulus processing

Our main finding is a long-lasting ERP effect of gain implicitly associated to inherently neutral faces that became evident from 200 to 700 ms after target face onset. Across the whole time window this ERP modulation consisted of increased centro-parietal positivities, presumably resembling P3 and LPC components - linked to higher-order stimulus evaluations - that were particularly boosted for gain-associated faces. Such modulations of late processing stages *(P3/LPC)* by monetary reward have been previously demonstrated in studies employing associative learning based on explicit valence categorization (Schacht et al. 2012; Rossi et al. 2017). These previous findings have been interpreted to indicate that previously rewarded stimuli receive increased cognitive resources, resulting in a prioritized processing (Nieuwenhuis et al. 2005), even for implicit reward associations (Bourgeois et al. 2016). In particular, the P3/LPC modulations on inherently neutral, but previously associated faces deserve special attention for two reasons: First, we did not find modulations of ERPs by motivational incentives after target face onset during the learning session. Second, the condition-to-face assignments were not made explicit for the participants during the learning session; indicating that the effect was driven by the implicit associations of the motivational contexts to certain faces. One potential explanation of these findings relates to the time required for consolidation that has been proposed in particular for arousing stimuli (Sharot et al. 2004). Therefore, overnight consolidation might play a crucial role particularly during the implicit association of motivational salience as similar P3 effects modulated by monetary reward were observed during an explicit learning paradigm without delay between learning and testing (Rossi et al. 2017).

In contrast to previous associative learning studies, in particular to Hammerschmidt and colleagues (Hammerschmidt et al. 2017) who detected P1 modulations driven by monetary reward associations, no ERP modulations at short latencies were found in the present study. Two reasons for this finding are conceivable: First, as early ERP effects of acquired salience were detected in studies employing explicit associative learning, implicitly learned associations might lead to less apparent impacts on perceptual encoding of the certain stimuli. Second, the task demands in the present study were exceptionally high and might have suppressed early ERP modulations (e.g., Pessoa 2014). In order to check whether the present study design actually allows for typical emotion-related ERP modulations, novel identities with facial expressions of emotion were presented in the same task during the test phase. Modulations of two emotion-related ERP components occurred: The face-sensitive N170 component was modulated by angry facial expressions compared to both neutral and happy expressions, supporting the assumption that the N170 is primarily (if at all) influenced by negative expressions (for reviews, see Rellecke et al. 2013; Hinojosa et al. 2015). It was further suggested that the N170 might be overlapped by the directly following EPN component which leads to comparable modulations by emotional expressions (Schacht and Sommer 2009; Rellecke et al. 2011; Rellecke et al. 2012). For the EPN component, typical modulations were found for happy and angry compared to neutral facial expressions (e.g., Hammerschmidt et al. 2017), as the EPN is known to reflect the automatic encoding of the emotional content of a given stimulus independent of task demands (Rellecke et al. 2011). In addition to N170 and EPN, previous studies reported even earlier (P1) or later LPC modulations (e.g., Schupp et al. 2004; Rellecke et al. 2012; Hammerschmidt et al. 2017), but in the present study those modulations were absent, potentially due to the task-irrelevance of the expressed emotion. Therefore, the present study design indeed allows for typical emotion-related ERP modulations; however, P1 modulations, known to be task-dependent (Pratt et al. 2011; Rellecke et al. 2011), might therefore be suppressed by the high cognitive load of the task used in the present study.

### Motivational contexts boost subsequent processing of even task-irrelevant stimuli

Recent studies provided robust evidence for impacts of motivational context on target stimulus processing (e.g., Krebs and Woldorff 2017), interestingly taking place even before effects of spatial attention occur (Bayer et al. 2017). What has yet been largely neglected is the question whether the motivational salience of cue stimuli might lead to preferential processing similar to stimuli of varying emotional content, such as affective scenes or emotional expressions (cf., Anderson 2013). Using cue stimuli of identical shape that only differed in color (counterbalanced), allowed us to investigate potential ERP modulations through the cues’ meaning, by keeping visual features constant across conditions. Interestingly, we found increased ERP effects to gain- and loss-indicating cues that resembled typical ERP modulations driven by stimuli of emotional content across different domains, i.e. EPN and LPC effects (e.g., Schacht and Sommer 2009; Bayer and Schacht 2014). This impression was verified by topography comparisons between these ERP responses to the cues during the learning session and to EPN effects elicited by emotional expressions during the test session in the present study. Importantly, the first visually evoked ERP component after cue onset – the P1- did not differ as a function of the cues’ motivational salience. As cue stimuli in the present study were perceptually identical besides variation in three equiluminant colors, the lack of P1 effects indicate that previously reported P1 effects modulated by emotional valence (e.g., Pourtois et al. 2004; Rellecke et al. 2012) reflect rapid core-feature analysis under the precondition that these features are clearly discriminable (Fedota et al. 2012).

Impacts of motivational incentives were, importantly, not restricted to the processing of cues but extended to the subsequent processing of even task-irrelevant stimuli within trials of increased motivational salience during the learning session. These impacts, however, declined when the target face was presented. As studies using associative learning paradigms typically report stabilized associated effects on target processing, future research is needed to determine the emergence of those associated effects.

### Effects on pupil dilations

In the learning session, pupil dilations were enlarged for both gain- and loss-related contexts compared to neutral contexts. These findings indicate increased arousal or attention triggered by motivational incentives (Massar et al. 2016; Pulcu and Browning 2017). In the test session, although LPC modulations driven by reward associations were detected on the neural level during, pupil size did not differ as a function of associated motivational salience, indicating that physiological arousal only increases when motivational incentives are directly available. Furthermore, pupil size was also not impacted by facial expressions carrying inherent emotional salience (although these elicited EPN modulations on the neural level), contradicting previous findings (Kret et al. 2013) and thus indicating that impacts of emotional expressions might be suppressed by the cognitive load of the task and the consequential task-irrelevance of the expressed emotion.

### Impacts of monetary gain and loss under conditions of equalized outcomes

In contrast to recent studies, which typically linked incentives explicitly to successful learning, the present study design ensured equalized outcomes of monetary gain and loss, but nevertheless demonstrated a prioritized neural processing of gain over loss. The influential prospect theory in economic decision making (Kahneman and Tversky 1979; Tversky and Kahneman 1992) already suggested an asymmetric function of gains and losses – with a typically higher impact of losses than gains during risky choices. This asymmetry is potentially based on the activation of different brain areas (Trepel et al. 2005), especially during reinforcement learning tasks (Wächter et al. 2009; Kim et al. 2015). In contrast, visual selective attention studies revealed an advantage of gains over losses in the prioritized processing (for a review, see Chelazzi et al. 2013). Recently, a first explanation for these seemingly conflicting assumptions was proposed based on findings that gain-associated targets were processed faster than loss-associated targets (Chapman et al. 2013). The authors concluded that the inhibition necessary for loss aversion takes more time than the facilitated processing elicited by reward associations.

### Conclusion

The present findings demonstrate that motivational contexts impacted pupil dilation and led to an ongoing influence on the neural processing of subsequent visual stimuli (fixation cross, prime/mask) during the learning session, however, not persisting to the target faces. During the test session, implicitly associated motivational salience impacted the processing of neutral faces, reflected in an enhanced centro-parietal ERP modulation for previously gain-associated target faces. In contrast, target faces expressing emotions (happy, angry) modulated the typical emotion-related EPN component, whereas P1 and LPC modulations were suppressed presumably by high demanding task requirements. Together, this study provides new evidence that neural representations of neutral stimuli can acquire increased salience via implicit learning, with an advantage for gain over loss associations.

## Acknowledgments

The authors thank Anna-Maria Grimm and Rebecca Jacob for their contributions to the development of the study design and data collection, Florian Niefind and Kay Reimers for their technical support during experimental setup, and Benthe Kornrumpf and Anton Unakafov for providing codes for data analyses.

